# A subset of human choroid plexus epithelial cells exhibit mitochondrial eccentricity and distinct expression of the pigmentation-associated enzyme TYRP1

**DOI:** 10.1101/2024.12.05.627055

**Authors:** Adam M.R. Groh, Shobina Premachandran, Finn Creeggan, Moein Yaqubi, Stephanie Zandee, Alexandre Prat, Jack P. Antel, G.R. Wayne Moore, Jo Anne Stratton

## Abstract

For decades, ultrastructural evaluation of epithelial cells in diverse organ systems has demonstrated the existence of two subtypes identified by stark differences in cytoplasmic electron density – so-called light and dark epithelial cells. Choroid plexus (CP) epithelial cells are key regulators of CSF homeostasis and are one of many specialized epithelial linings that exhibit this bimodal phenotype. Despite longstanding acknowledgement, it has been difficult to assess the potential significance of adult human light and dark CP epithelial cells due to a lack of characterization beyond electron microscopy (EM). We present the first transcriptomic analysis of adult human CP epithelial cells and denote the existence of four epithelial subpopulations, one of which is defined by elevated expression of *TYRP1* – a melanocyte-associated tyrosine-related protein involved in cellular pigmentation and proliferation. *TYRP1*-high cells also downregulate genes related to cilia function (which is consistent with observations of dark cell identity in organoids) and upregulate genes associated with pathways related to cell cycling, stress, and iron regulation. Our data provide an explanation of the molecular underpinning of adult human light and dark cell identity and serve as a resource for investigations of epithelial heterogeneity in the CP and other organs where dark cells are found.

## Introduction

Epithelial-like cells are thought to originate early in Metazoan evolution from the *Porifera* (sea sponge). Sponges utilize these cells to seal and regulate the passage of solutes to and from their internal milieu.^1,2^ With continued adaptation, epithelial cells began anchoring themselves to a basement membrane,^3^ ultimately leading to the emergence of more intricate and compact arrangements of cell sheets that line body cavities/organs.^4,5^ Epithelial cells have also developed diverse functional capabilities depending on their location,^6^ such as the secretion of specialized enzymes in the gastrointestinal tract, mucous in the respiratory tract, hormones in endocrine glands, and cerebrospinal fluid (CSF) in the brain.^7–10^

Despite such a broad range of organ-specific epithelial cell functions, ultrastructural evaluation by electron microscopy (EM) has consistently revealed two primary subtypes that exist independent of region and disease – so-called light and dark epithelial cells. These two epithelial subtypes have been noted in mouse tracheal,^11^ bile duct,^12^ renal,^13^ and choroid plexus (CP) epithelium,^14^ in addition to human gall bladder,^15^ large intestine,^16^ mammary gland,^17^ eccrine sweat gland,^18^ and choroid plexus epithelium.^19,20^ Despite clear ultrastructural differences between light and dark cells observed by EM, the functional significance of these epithelial subtypes remains unclear across organ systems.

CP epithelial cells have emerged as key players in brain homeostasis, contributing not only to CSF production, but to regulation of the blood-CSF-brain barrier and immune cell trafficking in a variety of neurological disorders.^21,22^ Light and dark CP epithelial cells have been noted in a number of mammalian species, including humans,^21^ yet their molecular profile is not known. While it is possible that the proportion of dark cells might increase in response to brain pathology,^23–25^ dark CP epithelial cells appear to constitute 12% of all epithelial cells in the mouse CP by embryonic day 14, suggesting that they are also present in health.^26^ Indeed, early work has demonstrated the existence of light and dark epithelial cells in normotypic human choroid across the lifespan.^20^ Dark CP epithelial cells are thought to have increased numbers of ribosomes and rough endoplasmic reticulum, thinner microvilli, and fewer microtubule-associated proteins.^14,27^ A recent study was the first to capture four epithelial cell subpopulations via single cell RNA sequencing (scRNA-seq) of developmental CSF-producing human organoids.^28^ They hypothesized that the two most abundant clusters in their epithelial dataset were light and dark cells, based on the upregulation of mitochondrial-related genes and downregulation of cilia- and microtubule-related genes in what they proposed to be a dark cell cluster.^28^ Although these findings provide a promising look into the molecular differences defining developmental light and dark CP epithelial cells in a model system, the data do not fully corroborate other well-described ultrastructural differences that are known to define dark cells, such as increased ribosomal content and altered cytoplasmic electron density (i.e., pigmentation), and the authors do not define a clear marker protein to differentiate these cell populations. Furthermore, a single-cell level characterization using adult human brain tissue is lacking.

We validate and build upon previous observations of human light and dark CP epithelial cells by quantifying ultrastructural differences in mitochondrial number, size, and morphology, in addition to qualitatively noting altered ribosomal content and basement membrane structure. We then provide the first adult human brain scRNA-seq profiling of the CP, which allowed us to reveal four subpopulations of CP epithelial cells, three of which can be defined by a gradient of *TYRP1* expression – a melanocyte-associated gene that produces tyrosinase-related protein 1, which is involved in melanin synthesis and cellular proliferation. *TYRP1*^high^ cells also upregulated a variety of genes related to mitochondria, ribosomes, stress, detoxification, iron regulation, and cell cycle arrest, consistent with our ultrastructural analysis. IHC confirmed expression of TYRP1 in a similar proportion of CP epithelial cells as past observations of dark cell proportions.^26^ These results provide a multimodal characterization of light and dark CP epithelial cells in the adult human brain. Our transcriptomic epithelial characterization could serve as a tool for future investigations that seek to explore the functional significance of these cells in the brain and other organ systems.

## Method

This research received ethical approval by the McGill University Faculty of Medicine Institutional Review Board (A09-M72-22B). Tissues from the Centre Hospitalier de l’Université de Montréal (CHUM) were obtained by a Material Transfer Agreement between CHUM and McGill University with full ethical approval (BH07.001, Nagano 20.332-YP) and informed consent. A tissue sample was also obtained from The Netherlands Brain Bank (NBB), Netherlands Institute for Neuroscience, Amsterdam (open access: www.brainbank.nl), where all material has been collected from donors for or from whom a written informed consent for a brain autopsy and the use of the material and clinical information for research purposes had been obtained by the NBB.

### Tissue processing for electron microscopy

CP tissue used for EM was extracted at the CHUM from the brains of 2 patients with Amyotrophic Lateral Sclerosis (ALS) within 4 hours post-mortem and immediately drop-fixed in a solution of 2.5% glutaraldehyde, 2% paraformaldehyde, and 0.1M sodium cacodylate in double-distilled water (ddH_2_O) for 24-48 hours at 4 °C, before transferring to phosphate-buffered saline (PBS). Tissues were cycled into fresh PBS once per week at 4 °C until processing. Once transported to the Facility for Electron Microscopy Research (FEMR) at McGill University, tissues were transferred back to the glutaraldehyde-based fixative solution with 4% sucrose overnight at 4 °C, and washed 3x by aspiration with 0.1M cacodylate washing buffer. Samples were post-fixed with 1% aqueous osmium tetroxide plus 1.5% aqueous potassium ferrocyanide for 2 hours at 4 °C, followed by three washes with ddH_2_O. Samples were then dehydrated in a standard set of acetone baths of increasing concentration at room temperature (RT) and subsequently immersed in 1:1 Epon:acetone overnight, 2:1 Epon:acetone the next day, and 3:1 Epon:acetone the following night, all on a rotator at RT. Tissues were then infiltrated with 100% Epon for four hours at RT with tube caps off: two hours on a rotator, and 2 hours under a vacuum. Specimens were then embedded in moulds, left at RT for 1 hour in a fume hood, and polymerized in an oven at 60 °C for 48 hours. Once polymerized, samples were removed from the oven, cooled at RT, and sectioned. Sections were stained with toluidine blue to confirm tissue integrity. Separate 100nm-thick counter-stained sections were placed on TEM grids for imaging.

### Single cell RNA sequencing

Human CP samples were obtained from patients with ALS (n=3) and Hereditary Spastic Paraplegia (n=1) who underwent medical assistance in dying at CHUM. CP tissue samples (n=4) used for scRNA-seq were extracted from the brain within 4 hours post-mortem, dissociated, and cell suspensions loaded into a 10x Chromium machine for GEM generation. Libraries were prepared as per manufacturer’s recommendations and sequenced. Fastq files were generated and used as input into the cellranger software, which generated filtered count matrices that were imported in R (v.4.2.3) using the Seurat (v4.3.0.1) package. Datasets were preprocessed, normalized and integrated using standard Seurat functions. PCA, UMAP, and gene expression calculations were also conducted using Seurat.

### Immunohistochemistry (IHC)

CP tissue used for IHC was extracted at the CHUM from the brain within 4 hours post-mortem and drop-fixed in 10% neutral buffered formalin. Frozen CP tissue from the Netherlands Brain Bank from a patient with major depressive disorder (MDD) was fixed in 10% neutral buffered formalin and processed via a standard set of alcohol and xylene incubations. Dehydrated samples were embedded in paraffin, sectioned (15 μM), and mounted. Formalin-fixed paraffin-embedded (FFPE) sections were dewaxed with xylene, rehydrated in a series of ethanol solutions, and rinsed in ddH_2_O. Antigen retrieval was conducted by immersing slides in citrate buffer (pH 6; in PBS) in an antigen retrieval chamber for 10 min at 95 °C. Slides were left to cool at RT and washed in PBS. Blocking was performed for 60 minutes at RT with goat blocking buffer from the Tyramide SuperBoost Kit (B40922, B40923, Invitrogen). Primary antibodies were diluted in 2% BSA in PBS and incubated overnight at 4 °C. These included TYRP1 (rabbit; 1:200; PA5-81909, Invitrogen) and vimentin (mouse; 1:200; c9080, Sigma). Poly-HRP conjugated secondary antibodies were then incubated for 60 minutes at RT followed by 3x PBS washes. 100µl tyramide working solution was incubated for 8min at RT, followed by 100 µl of stop reagent for 5 min, and 3x washes in PBS. DAPI (1:1000 in PBS; D1306, Invitrogen) was applied for 5 min followed by 3x washes in PBS. Prolong Gold Antifade Mounting Medium (P10144, Invitrogen) was used to mount.

### Confocal and electron microscopy

Fluorescent images of CP epithelial cell antigens used for quantification were taken with the same acquisition parameters on a Leica TCS SP8 multiphoton confocal microscope at 40x or 63x magnification in oil with additional optical zoom. Representative images were obtained using the HyVolution setting on the Leica TCS SP8 at 63x, with additional optical zoom. Cell counts were completed using Fiji. TEM grids were imaged on a FEI Tecnai G2 Spirit Twin 120 kV Cryo-TEM. Mitochondria in light and dark CP epithelial cells (n=17 light, n=17 dark) were counted and normalized to the area of the cell size (in pixels) using Fiji. Five mitochondria in each cell used for mitochondrial count analysis were randomly selected, manually segmented, and circularity was determined using the shape measurement function (1.0 = perfectly circular) in Fiji. All appropriate statistical tests were run using GraphPad Prism (v8.0.1).

## Results

### Human dark CP epithelial cells possess increased numbers of eccentric mitochondria, elevated mitochondrial and cytoplasmic electron density, and basement membrane convolutions

It has been observed that mammalian dark CP epithelial cells possess greater ribosomal content, increased cytoplasmic, nuclear, and microvilli electron density, and elevated levels of intracytoplasmic osmiophilic droplets; some have speculated that the general dark quality of these cells could be explained by an altered hydration state at time of fixation, although this has never been confirmed.^19,29^ Recent work in developmental human organoids noted increased numbers of mitochondria in dark cells.^28^ Our ultrastructural analysis confirmed the presence of dark cells in the adult human CP (Figure 1A). We also corroborate that dark CP epithelial cells in the adult human brain possess increased numbers of mitochondria (p<0.01; Figure 1B and C). In addition, we note that dark cell mitochondria are highly eccentric, in that they deviate from a perfect circular shape (Figure 1B). Light CP epithelial cell mitochondria had an average circularity score of 0.920 (with 1 indicating perfect circularity), whereas dark CP epithelial cell mitochondria had an average score of 0.748 (p<0.0001; Figure 1C). Dark cell mitochondria were also smaller (p<0.0001; Figure 1C) and more electron dense (p<0.0001; Figure 1C). The cytoplasm of dark CP epithelial cells was also substantially more electron dense (p<0.0001; Figure 1C). Further qualitative examination of dark CP epithelial cells suggested that their basement membranes are convoluted compared to light cells (Figure 1B; Supplemental Figure 1) and that they possess more cytoplasmic polyribosomes (Figure 1B). Together, these data demonstrate stark ultrastructural differences between light and dark CP epithelial cells that likely have implications for cell function.

**Figure 1.**
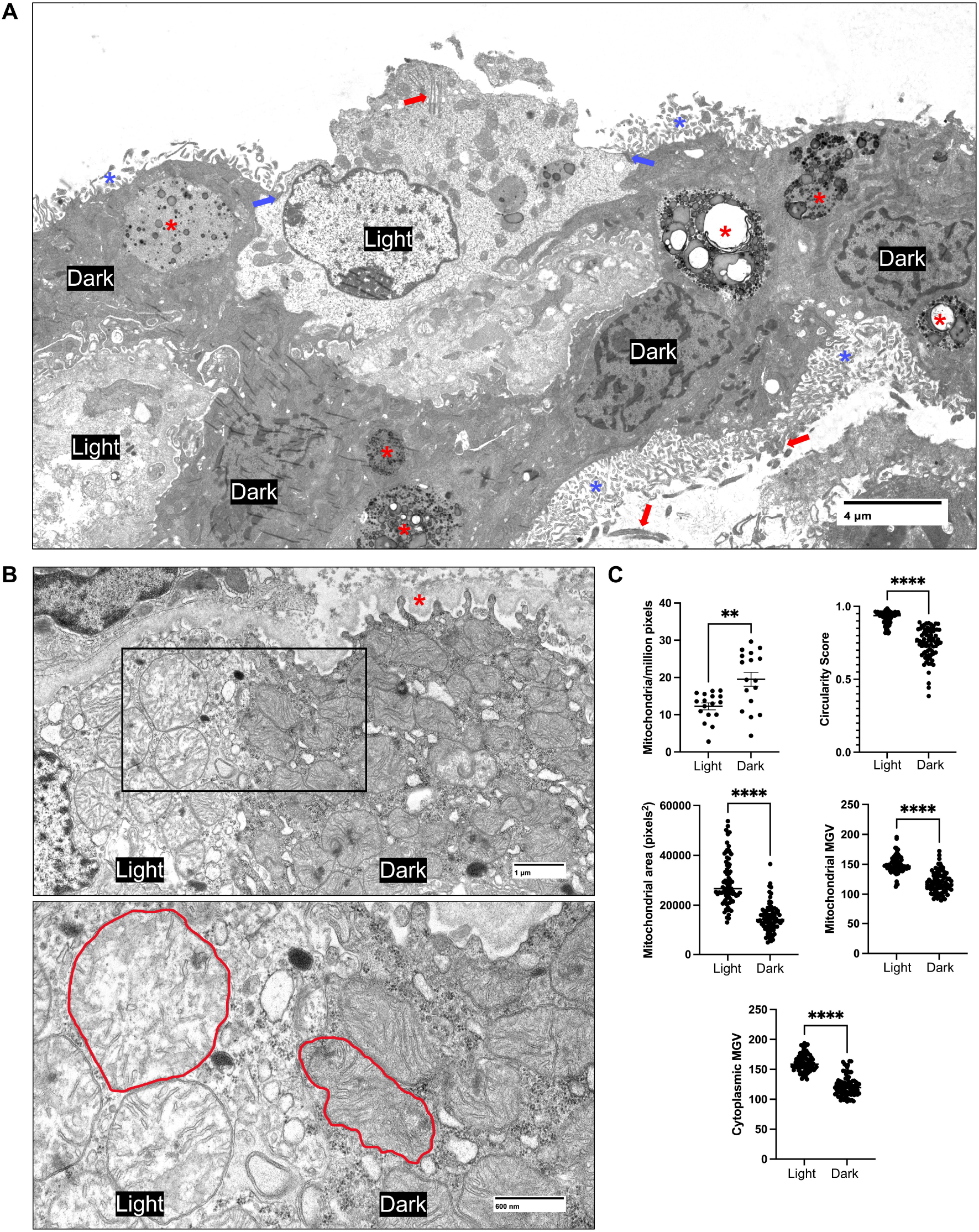
Human dark CP epithelial cells possess increased numbers of eccentric mitochondria, elevated mitochondrial and cytoplasmic electron density, and abnormal basement membrane convolutions. (A) Low magnification image of a collection of dark CP epithelial cells surrounding a light CP epithelial cell. Dark cells appear to be arranged in patches, rather than scattered evenly throughout the choroid epithelium. Red asterisks indicated the presence of lipofuscin (granules of lysosomal debris containing lipid), which appears to be elevated in dark CP epithelial cells. Blue asterisks indicate the location of apical microvilli, red arrows denote cilia, and blue arrows indicate apical tight junctions. Each CP epithelial cell type is labeled as “light” or “dark”, respectively. Scale bar = 4 µm. (B) Magnified image of a light-dark CP epithelial cell transition. The red asterisk indicates a convoluted basement membrane attaching the dark CP epithelial cell to the underlying choroid stroma, which immediately transitions to a flat basement membrane anchoring the light epithelial cell. A black box indicates the location of the magnified image below, which is used to demonstrate the differences in mitochondrial structure between light and dark CP epithelial cells. In the magnified image, red lines outline representative examples of mitochondrial morphology in light and dark CP epithelial cells; 2D sections of light cell mitochondria are rounded and typically larger, whereas sections of dark cell mitochondria are smaller and highly eccentric (deviating from a circular shape). A general increase in electron density of dark CP epithelial cell cytoplasm and mitochondria can also be observed in both images in panel B, in addition to an increased presence of polyribosomes. Each CP epithelial cell type is labeled as “light” or “dark”, respectively. Scale bars = 1 µm and 600 nm. (C) Quantification of some ultrastructural observations in light and dark CP epithelial cells: the average number of mitochondria (per million pixels), the circularity of mitochondria, the area of mitochondria (pixels^2^), and the mitochondrial and cytoplasmic electron density (mean gray value [MGV]).

### Single cell RNA sequencing of the adult human CP suggests the presence of four distinct epithelial cell subpopulations

To characterize the epithelial cells of the human CP, we profiled 4030 cells from this region. Using the expression of canonical cell type markers, we identified several distinct cell types: endothelial cells, epithelial cells, immune cells, fibroblasts, pericytes, and red blood cells (Figure 2A). Re-clustering of 395 epithelial cells expressing the canonical markers *TTR* and *ENPP2* (Figure 2B) revealed four epithelial cell clusters. We defined these four populations as TYRP1-high, TYRP1-intermediate, TYRP1-low, and FABP7-high epithelial cells based on differential gene expression analysis (Figure 2C-E). Trajectory analysis indicated that TYRP1-high and TYRP1-low epithelial cells occupy opposite ends of a developmental trajectory, with the FABP7-high epithelial cell population positioned at an intermediate node (Figure 2F). FABP7-high epithelial cells were then excluded from downstream analysis to permit further characterization of TYRP1-high and low epithelial cells. We observed that *TYRP1* exhibited a gradient of expression across the three remaining clusters (Figure 2G). Gene ontology analysis of upregulated genes in each CP epithelial cluster revealed distinct biological processes indicative of CP epithelial subtype identity. TYRP1-high cells upregulated genes associated with inclusion body assembly and apoptotic mitochondrial changes (Figure 2H); these terms corresponded well to the altered mitochondrial morphology (Figure 1B) and considerable lipofuscin and osmophilic granule buildup we observed in dark cells (Figure 1A; Supp. Figure 1). Conversely, TYRP1-low epithelial cells upregulated genes associated with cilium assembly, basement membrane organization, and regulation of bilateral symmetry (Figure 2H), strongly suggesting that these cells were light cells based on previous descriptions in organoids.^28^ TYRP1-intermediate epithelial cells upregulated genes associated with cellular respiration, and shared upregulation of genes associated with ribosome biogenesis and apoptosis with TYRP1-high cells (Figure 2H), suggesting they might represent an epithelial subtype in transition from light to dark. Further analysis of the TYRP1-high (dark cell) cluster suggested strong involvement in pigment granule generation, cell cycling, proliferation, iron binding, and apoptotic regulation (Figure 2I). When we compiled a list of genes implicated in melanin production, iron sequestration, extracellular matrix organization, and cell cycle arrest, we found that TYRP1-high epithelial cells had the highest expression levels of these genes, followed by TYRP1-intermediate (stressed) and TYRP1-low (light) cells (Figure 2J), mirroring the expression pattern of *TYRP1* (Figure 2G). In summary, we provide the first transcriptomic characterization of adult human TYRP1-high (dark) and TYRP1-low (light) EP epithelial cells.

**Figure 2.**
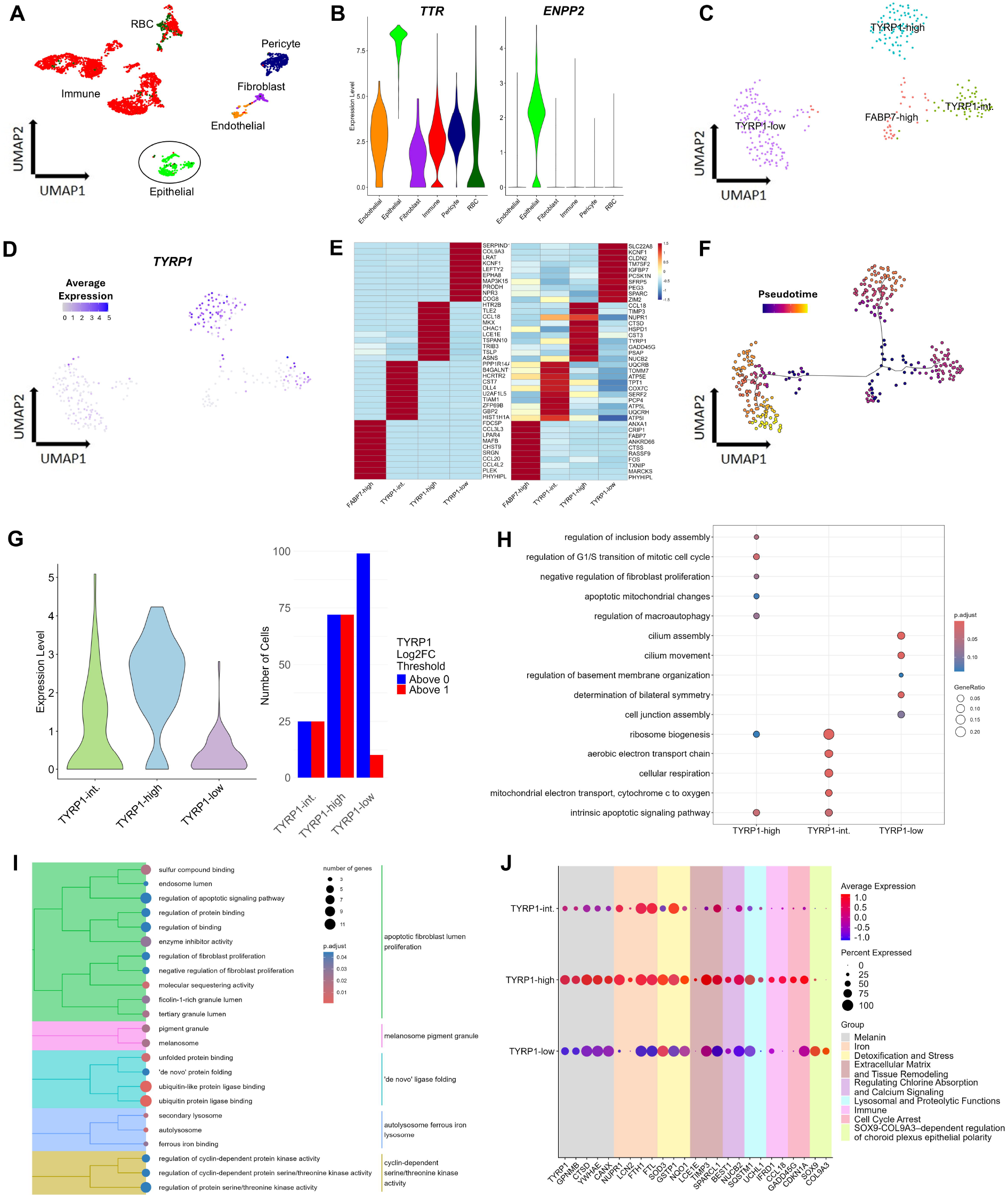
Single cell RNA sequencing of adult human choroid plexus suggests the presence of four distinct epithelial cell subpopulations. (A) UMAP plot depicting the major CP cell types identified by scRNA-seq. Epithelial cells are circled in black. (B) Violin plot showing the expression of epithelial-specific genes *TTR* and *ENPP2* across the identified cell types. (C) UMAP plot displaying four identified epithelial subtypes. (D) FeaturePlot displaying expression of the *TYRP1* gene epithelial subtypes. (E) Heatmap showing the expression profiles of the top 10 differentially upregulated genes per cluster, based on fold change (left panel) and adjusted p-value (right panel). The average expression of genes is represented by a color gradient, with red indicating high expression and blue indicating low expression. (F) Pseudotime plot illustrating the trajectory of the identified epithelial subtypes. Pseudotime score is represented using a color gradient. (G) Violin plot visualizing expression of *TYRP1* across TYRP1-high, TYRP1-intermediate, and TYRP1-low epithelial cell subtypes (left panel) and a bar graph demonstrating expression of *TYRP1* using two different thresholds. Cells expressing any level of *TYRP1* are shown in blue, and cells with *TYRP1* expression greater than 1 (based on log2 fold change) are shown in red. (H) DotPlot illustrating selected biological processes unique to TYRP1-high, TYRP1-intermediate, and TYRP1-low epithelial subtypes. Two shared biological processes between TYRP1-high and TYRP1-intermediate epithelial subtypes are also included. The gene ratio for each biological term is represented by the size of the circles, and statistical significance is indicated by a color gradient, with red showing high significance and blue showing low significance. (I) Tree plot of the most upregulated biological processes in TYRP1-high cells compared to other epithelial subtypes. The number of genes associated with each term is represented by circle size, and statistical significance is indicated by a color gradient, with red showing high significance and blue showing low significance. (J) DotPlot showing the expression of compiled marker genes across the three epithelial subtypes. The percentage of cells expressing each gene is represented by circle size, and genes associated with different biological processes are colored distinctly. The name of each gene group is shown in the legend to the right of the plot. The average expression of genes is represented by a color gradient, with red indicating high expression and blue indicating low expression.

### Immunohistochemical analysis confirms the existence of a subset of TYRP1-high CP epithelial cells in the adult human brain that resemble dark cells

The first and only identification of human dark CP epithelial cells using IHC was completed in developmental human organoids by demonstrating increased expression of the mitochondrial protein CARD19 and decreased expression of the cilia transcription factor FOXJ1 in the same population of CP epithelial cells.^28^ We demonstrate that, beyond the long-described mitochondrial phenotype defining dark CP epithelial cells, they can also be identified by elevated expression of TYRP1 (Figure 3A and Bi-iii). Interestingly, 12-20% of all epithelial cells in the CP were TYRP1-high (p<0.0001), suggesting conservation of a similar proportion of dark CP epithelial cells between humans and mice.^26^ Further, we demonstrate that these cells exist in a similar proportion in the brains of patients with MS, ALS, and major depressive disorder, suggesting they are maintained at similar levels across diseases, and might have a function in health (Figure 3Biv).

**Figure 3.**
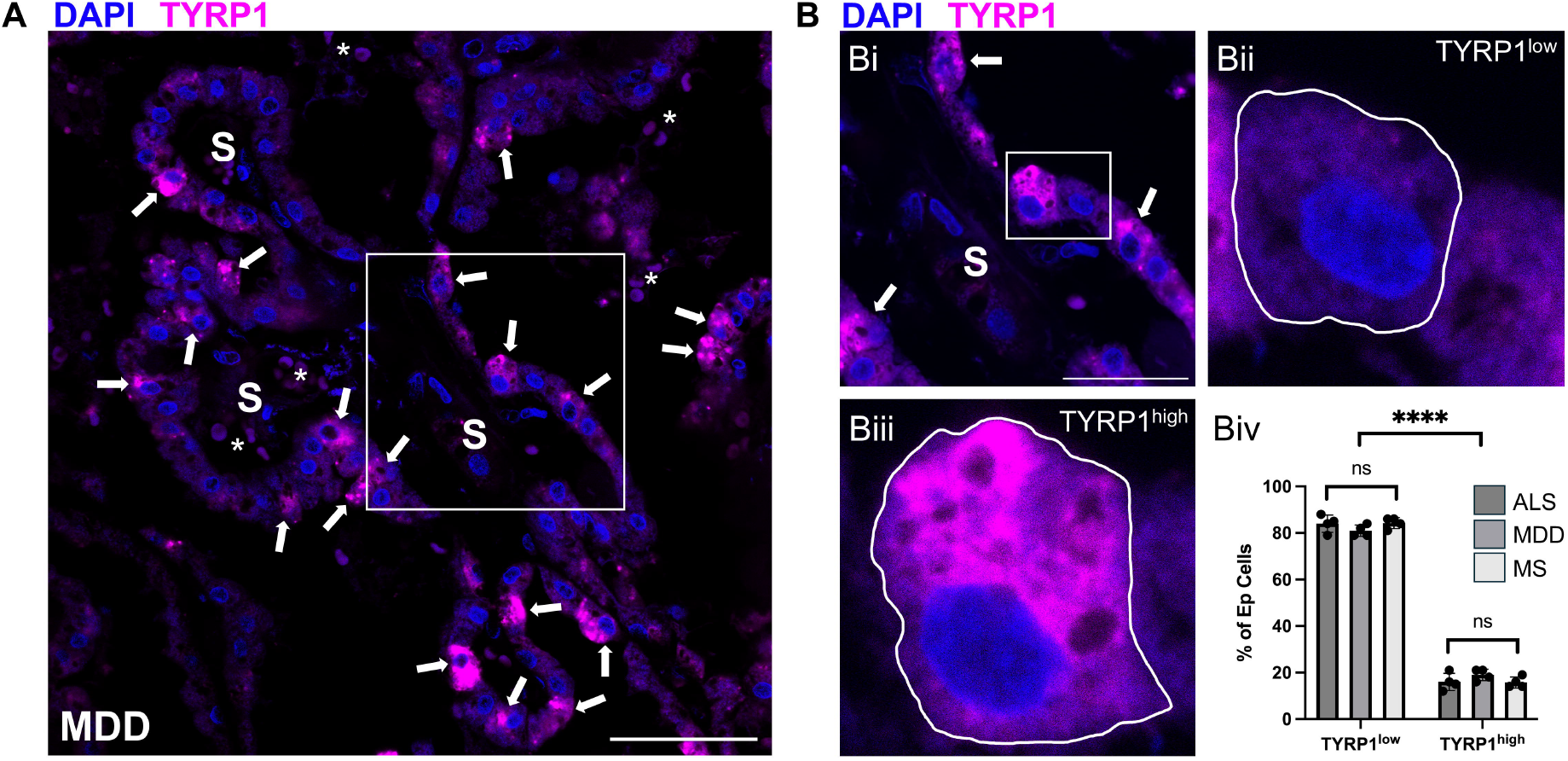
Immunohistochemical analysis confirms the existence of a subset of TYRP1-high CP epithelial cells in the adult human brain that resemble dark cells. (A) Low magnification image of TYRP1-high CP epithelial cells in a patient with major depressive disorder (MDD). White arrows denote TYRP1-high epithelial cells, and white asterisks denote autofluorescent red blood cells present in the fenestrated blood vessels of the choroid stroma. A white box indicates the location of cells magnified in panel B. S = stroma. Scale bar = 50uM. (B) (Bi) Higher magnification image depicting TYRP1-high and TYRP1-low CP epithelial cells. A white box indicates the location of cells magnified in panels Bii and Biii. (Bii) Magnified image of a single TYRP1-low CP epithelial cell. (Biii) Magnified image of a single TYRP1-high CP epithelial cell. (Biv) Quantification of the percentage of TYRP1-high and TYRP1-low CP epithelial cells in 1 ALS patient, 1 MDD patient, and 1 MS patient. No significant differences in TYRP1-high (dark) and TYRP1-low (light) CP epithelial cell proportions between conditions were found. TYRP1-high CP epithelial cells makeup approximately 15-20% of all epithelial cells (p<0.0001), similar to past observations of dark cells in rodents.

## Discussion

Dark CP epithelial cells in the adult human brain have been acknowledged for decades but their relevance to putative CP functions remains elusive.^21^ This is largely because they have been identified solely by electron microscopy with no way to selectively assess their function *in vivo*. By conducting whole cell scRNA-seq and EM on high-quality human CP tissue sourced from rapid autopsy, we provide a comprehensive description of CP epithelial cell subtypes in the adult human brain. We demonstrate that beyond previously-defined cilia and mitochondria-related gene expression differences uncovered in organoid model systems, dark CP epithelial cells are also defined by prominent upregulation of the melanocyte-associated *TYRP1*, in addition to a variety of other genes related to pigmentation, iron regulation, and stress. Together, these data provide a comprehensive transcriptomic characterization of light and dark CP epithelial cells and suggest that dark cells may be in a stressed or dysfunctional state which appears to be less amenable to maintenance of CSF homeostasis, based on their genetic profile.

One of the primary differentially-expressed genes we uncovered in dark CP epithelial cells was *TYRP1*. In animal models, *TYRP1* is critical for the synthesis of eumelanin, and has also been shown to play a non-enzymatic role in pigmentation via regulation of TYR stability.^30^ Beyond pigmentation, *TYRP1* mRNA has also been shown to ward off apoptosis in melanocyte cell lines. Targeting *TYRP1* with antisense oligonucleotides in melanocytes leads to G1 arrest and increased apoptosis *in vitro*.^31^ It is possible that *TYRP1* upregulation in dark CP epithelial cells represents a compensatory mechanism at play to overcome stress. This hypothesis is supported by concomitant upregulation of a variety of other genes involved in the cellular stress response and detoxification (*SOD3, GSTP1, NQO1*), cell cycle arrest (*GADD45G, CDKN1A*) and iron regulation/ferroptosis (*NUPR1, LCN2, FTH1*). Further, mitochondrial shrinking is a feature of ferroptosis,^32^ and we identified increased numbers of irregularly-shaped, smaller mitochondria in dark CP cells compared to light CP cells. Taken together, both scRNA-seq and ultrastructural data suggest that human dark CP epithelial cells are in a stressed state.

Given dark CP epithelial cells appear to be considerably more stressed than light CP epithelial cells, we speculated that this might affect putative CP functions. Indeed, we also noted that dark CP epithelial cells upregulate *BEST1*, a calcium-activated chloride channel. Intrathecal administration of arginine vasopressin has been shown to increase the number of dark CP epithelial cells in rats and reduce chloride ion efflux and CSF formation, leading the authors to suggest that dark CP epithelial cells modify fluid transfer.^23,33^ It is possible that the *BEST1* upregulation we observed in dark CP epithelial cells may be linked to altered chloride efflux, which could alter CSF secretion. Another study in rodents noted that dark CP epithelial cells resemble a “fluid absorbing structure”.^34^ The dominant presence of dark cells in preclinical models of hydrocephalus and other conditions of water-imbalance lends support for a potential role in reabsorption,^25^ as does the fact that an increase in light cell proportion is associated with a secretory state in hydrocephalic-polydactyl (hpy/hpy) mice.^35^

We also noted prominent convolutions in the basement membrane of dark cells compared to light cells – a cellular alteration which could also indicate dysregulated secretory capacity and/or altered epithelial polarity and cell-cell adhesion. In a recent investigation, it was found that *SOX9* is required for transcriptional upregulation of *COL9A3* in CP epithelial cells; loss of collagen IX led to severe disruption of the basement membrane, extracellular matrix dysregulation, altered apicobasal epithelial cell polarity, and ultimately a hyperpermeable blood-CSF-brain barrier.^36^ Strikingly, both *SOX9* and *COL9A3* were significantly downregulated in dark cells in our study, providing a potential genetic underpinning for the altered basement membrane phenotype we observed by EM, and suggesting that dark cells may contribute to changes in blood-CSF-brain barrier permeability. Recent work demonstrates that a subset of CP epithelial cells transiently specialize to coordinate immune cell recruitment and differentiation in response to LPS exposure, while also increasing the permeability of the BCSFB^22^; it is possible that dark cells represent an epithelial subpopulation that is more predisposed to entering this reactive state, although this requires further experimentation.

In summary, we complete the first adult human brain characterisation of light and dark CP epithelial cells and provide mechanistic hypotheses about cell function. Our results serve as a resource for future investigations of this curious epithelial subtype in the CP, and perhaps in other organ systems where light and dark epithelial cells are found.

## Data availability

Our scRNA-seq dataset has been uploaded to our open science website <u>SingloCell</u> so that other groups can explore the data further.

## Acknowledgements

The authors would like to thank the Facility for Electron Microscopy Research (<u>FEMR)</u> at McGill University for their support of this work. In particular, they extend gratitude to Kelly Sears for his insight into the operation of all instruments at the FEMR, and to Johanne Ouellette for meticulously processing and sectioning all tissue utilized for EM. Finally, they offer their sincere appreciation to the donors who made the research possible.

## Supplementary material

A single supplemental figure has been provided.

## Figure Legends

**Supplemental Figure 1**. Additional low magnification image of a single dark and light epithelial cell and underlying choroid stroma. Red asterisks indicate areas of convoluted basement membrane present underneath the dark cell. Blue arrows denote intact apical tight junctions, the red arrow points to a collection of cilia in cross-section, and the white asterisk demarcates a large osmophilic granule in the dark cell. Each CP epithelial cell type is labeled as “light” or “dark”, respectively. Scale bar = 2 µm.

## References

1. Leys SP, Nichols SA, Adams EDM. Epithelia and integration in sponges. Integr Comp Biol. 2009;49(2):167–177. doi:10.1093/icb/icp038

2. Adams EDM, Goss GG, Leys SP. Freshwater sponges have functional, sealing epithelia with high transepithelial resistance and negative transepithelial potential. PLoS One. 2010;5(11). doi:10.1371/journal.pone.0015040

3. Tyler S. Epithelium-The Primary Building Block for Metazoan Complexity 1. Vol 43.; 2003. https://academic.oup.com/icb/article/43/1/55/604456

4. Clemente-Suárez VJ, Martín-Rodríguez A, Redondo-Flórez L, Villanueva-Tobaldo CV, Yáñez-Sepúlveda R, Tornero-Aguilera JF. Epithelial Transport in Disease: An Overview of Pathophysiology and Treatment. Cells. 2023;12(20). doi:10.3390/cells12202455

5. Muse ME, Shumway KR, Crane JS. Physiology, Epithelialization. StatPearls Publishing; 2023.

6. Lowe JS, Anderson PG. Stevens & Lowe’s Human Histology, Epithelial Cells. 4th ed. Mosby; 2015.

7. Kurosumi K. Electron Microscopy in Biology and Medicine, Ultrastructure of Endocrine Cells and Tissues, Mechanism of Secretion in Endocrine Glands. Vol 1. Springer; 1984.

8. Terada T, Kitamura Y, Ashida K, et al. Expression of pancreatic digestive enzymes in normal and pathologic epithelial cells of the human gastrointestinal system. Virchows Archiv: European Journal of Pathology. Published online 1997:195-203.

9. Adler KB, Tuvim MJ, Dickey BF. Regulated mucin secretion from airway epithelial cells. Front Endocrinol (Lausanne). 2013;4(SEP). doi:10.3389/fendo.2013.00129

10. Damkier HH, Brown PD, Praetorius J. Cerebrospinal fluid secretion by the choroid plexus. Physiol Rev. 2013;93(4):1847–1892. doi:10.1152/physrev.00004.2013

11. Hansell MM, Moretti RL. Ultrastructure of the mouse tracheal epithelium. J Morphol. 1969;128(2):159–169. doi:10.1002/jmor.1051280203

12. Yamada K. Fine Structure of Rodent Common Bile Duct Epithelium. Vol 105.; 1969.

13. Clark SL. Cellular Differentiation in the Kidneys of Newborn Mice Studied with the Electron Microscope. J Biophys Biochem Cytol. 1957;3(3):349–362. doi:10.1083/jcb.3.3.349

14. Sturrock RR. A Morphological Study of the Development of the Mouse Choroid Plexus. Vol 129.; 1979.

15. Gilloteaux J, Pomerants B, Kelly TR. Human Gallbladder Mucosa Ultrastructure: Evidence of Lntraepithelial Nerve Structures. Vol 184.;. Vol 1989.

16. Shamsuddin AM, Phelps PC, Trump BF. Human Large Intestinal Epithelium: Light Microscopy, Histochemistry, and Ultrastructure. Hum Pathol. 1982;13(9):790–803. doi:10.1016/s0046-8177(82)80075-0

17. KellokumpulLehtinen P, Johansson R lM, Pelliniemi LJ. Ultrastructure of human fetal mammary gland. Anat Rec. 1987;218(1):66–72. doi:10.1002/ar.1092180111

18. Bovell D. The human eccrine sweat gland: Structure, function and disorders. Journal of Local and Global Health Science. 2015;2015(1). doi:10.5339/jlghs.2015.5

19. Dohrmann GJ, Bucy PC. Human choroid plexus: a light and electron microscopic study. J Neurosurg. 1970;(5):506–516. doi:10.3171/jns.1970.33.5.0506

20. Dohrmann GJ. Dark and Light Epithelial Cells in the Choroid Plexus of Mammals. Journal of Ultrastructural Research. 1970;32:268–273. doi:10.1016/S0022-5320(70)80007-7

21. Saunders NR, Dziegielewska KM, Fame RM, Lehtinen MK, Liddelow SA. The choroid plexus: a missing link in our understanding of brain development and function. Physiol Rev. 2023;103(1):919–956. doi:10.1152/physrev.00060.2021

22. Xu H, Lotfy P, Gelb S, et al. The choroid plexus synergizes with immune cells during neuroinflammation. Cell. Published online 2024. doi:10.1016/j.cell.2024.07.002

23. Johanson CE, Preston JE, Chodobski A, et al. AVP V 1 Receptor-Mediated Decrease in Cl Efflux and Increase in Dark Cell Number in Choroid Plexus Epithelium.; 1999.

24. Engelhardt B, Wolburg-Buchholz K, Wolburg H. Involvement of the Choroid Plexus in Central Nervous System Inflammation. Microsc Res Tech. 2001;52(1):112–129. doi:10.1002/1097-0029(20010101)52:1<112::AID-JEMT13>3.0.CO;2-5

25. Weaver CE, McMillan PN, Duncan JA, Stopa EG, Johanson CE. Hydrocephalus disorders: their biophysical and neuroendocrine impact on the choroid plexus epithelium. Advances in Molecular and Cell Biology. 2004;31:269–293.

26. Wolburg H, Paulus W. Choroid plexus: Biology and pathology. Acta Neuropathol. 2010;119(1):75–88. doi:10.1007/s00401-009-0627-8

27. Pellegrini L, Lancaster MA. Breaking the barrier: In vitro models to study choroid plexus development. Curr Opin Cell Biol. 2021;73:41–49. doi:10.1016/j.ceb.2021.05.005

28. Pellegrini L, Bonfio C, Chadwick J, Begum F, Skehel M, Lancaster MA. Human CNS barrier-forming organoids with cerebrospinal fluid production. Science (1979). 2020;369(6500). doi:10.1126/science.aaz5626

29. Wislocki GB, Ladman AJ. The Fine Structure of the Mammalian Choroid Plexus. (Wolstenholme G, O’Connor CM, eds.). Wiley; 1958.

30. Gautron A, Migault M, Bachelot L, Corre S, Galibert MD, Gilot D. Human TYRP1: Two functions for a single gene? Pigment Cell Melanoma Res. 2021;34(5):836–852. doi:10.1111/pcmr.12951

31. Li CY, Gao TW, Wang G, et al. The effect of antisense tyrosinase-related protein 1 on melanocytes and malignant melanoma cells. British Journal of Dermatology. 2004;150(6):1081–1090. doi:10.1111/j.1365-2133.2004.05929.x

32. Li J, Cao F, Yin H liang, et al. Ferroptosis: past, present and future. Cell Death Dis. 2020;11(2). doi:10.1038/s41419-020-2298-2

33. Liszczak TM, Black 1’ ZPM, Foley L. Cell and Tissue Arginine Vasopressin Causes Morphological Changes Suggestive of Fluid Transport in Rat Choroid Plexus Epithelium. Vol 246.; 1986.

34. Schultz WJ, Brownfield MS, Kozlowski GP. Cell and Tissue Research The Hypothalamo-Choroidal Tract II. Ultrastructural Response of the Choroid Plexus to Vasopressin *. Vol 178.; 1977.

35. Shuman C, Bryan J. Comparative quantitative ultrastructural studies of the choroidal epithelium of hydrocephalic (hpy/hpy) and normal mice, and the effect of stress induced by water deprivation. Anat Anz. 1991;173(1):33–44.

36. Vong KI, Ma C, Li B, et al. SOX9-COL9A3-dependent regulation of choroid plexus epithelial polarity governs blood-cerebrospinal fluid barrier integrity. Proc Natl Acad Sci U S A. 2021;118(6). doi:10.1073/pnas.2009568118/-/DCSupplemental

