## Supplementary figures and images for "A subset of human choroid plexus epithelial cells exhibit mitochondrial eccentricity and distinct expression of the pigmentation-associated enzyme TYRP1"

### Supplemental Figure 1

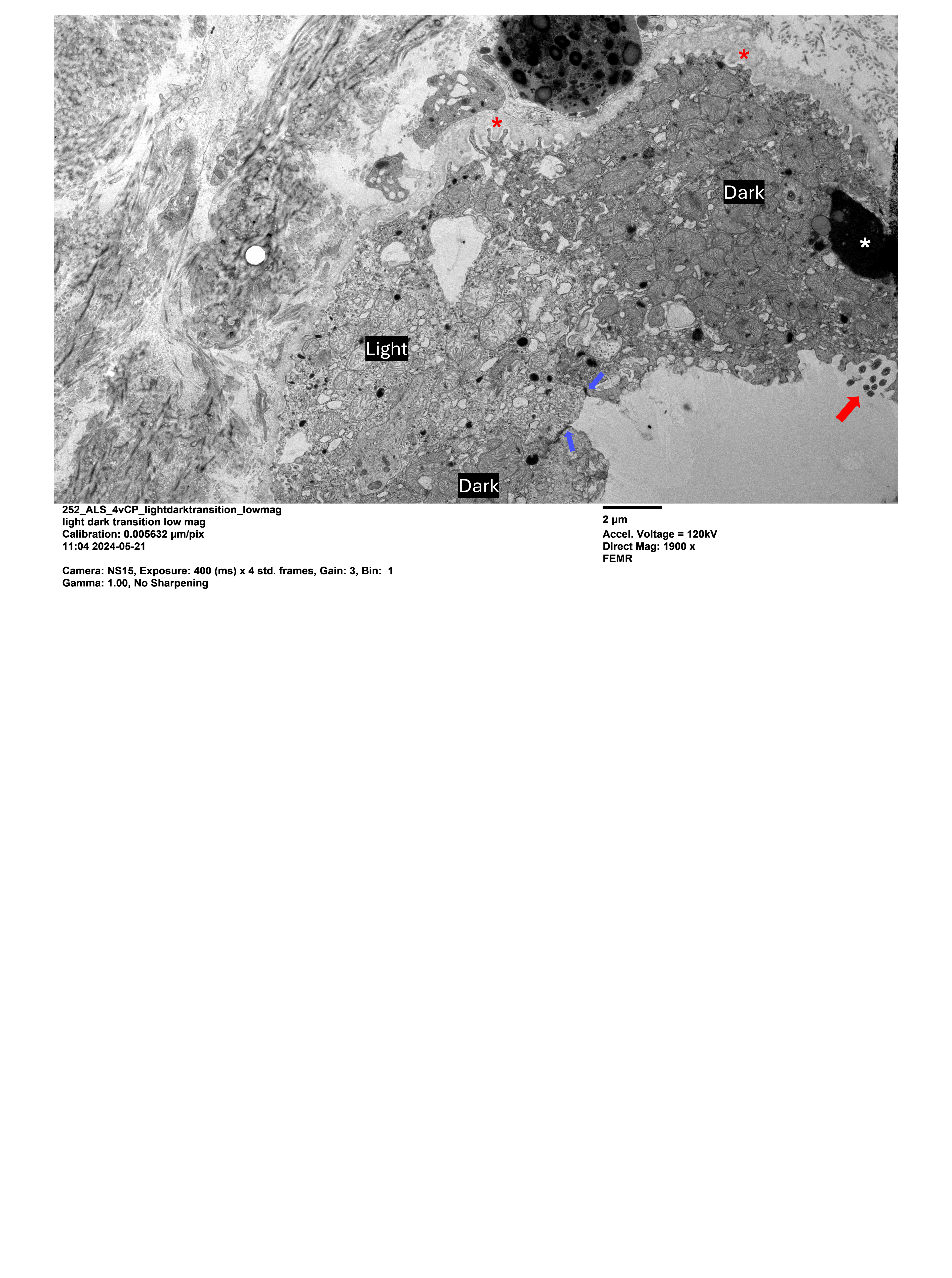
